# Humans use minimum cost movements in a whole-body task

**DOI:** 10.1101/2020.12.29.424756

**Authors:** Lijia Liu, Dana Ballard, Mary Hayhoe

**Affiliations:** Department of Computer Science, The University of Texas at Austin, Austin, TX; Center for Perceptual Systems, The University of Texas at Austin, Austin, TX

**Keywords:** Posture analysis, whole body movement, virtual tracing, kinematic representation, movement variation costs

## Abstract

Humans have elegant bodies that allow gymnastics, piano playing, and tool use, but understanding how they do this in detail is difficult because their musculoskeletal systems are extraordinarily complicated. Nonetheless, common movements like walking and reaching can be stereotypical, and a very large number of studies have shown their movement cost a major factor. In contrast, one might think that general movements are very individuated and intractable, but a recent study has shown that in an arbitrary set of whole-body movements used to trace large-scale closed curves, near-identical posture sequences were chosen across different subjects, both in the average trajectories of the body’s limbs and in the variance within trajectories. The commonalities in that result motivate explanations for its generality. One possibility could be that humans also choose trajectories that are economical in energetic cost. To test this hypothesis, we situate the tracing data within a fifty degree of freedom dynamic model of the human skeleton that allows the computation of movement cost. Comparing the model movement cost data from nominal tracings against various perturbed tracings shows that the latter are more energetically expensive, inferring that the original traces were chosen on the basis of minimum cost.

## 1. Introduction

A general principle of human movement is that our nervous system should exhibit trajectories that are economical in energetic cost [1, 2]. It has been established for decades and has been well studied in simple movements. In locomotion, there are a number of experiments showing that humans’ walking speed [3], step frequency/length [4, 5, 6, 7, 8, 9, 10], step width [11, 12] are all corIn particularrelated with the minimum metabolic cost, In particular, energetic cost exhibits a U-shaped dependence on step frequency while walking at a constant speed [13, 8], and the minimum of the U-shape is consistent with the self-selected or preferred walking frequency. Furthermore, new evidence [14, 15, 16] shows the system can adapt preferred gaits to minimize energetic cost in response to varying loads.

Although the principle that humans’ self-selected trajectories or posture sequences are economical in energetic cost has been commonly shown in the studies of simple single-behavior motions such as walking, running, and reaching, whether the principle is true for large-scale complex movements still needs to be tested. Thus, we conducted a complex whole-body virtual tracing experiment [17] that aimed to learn the principles behind *large-scale arbitrary* movements, particularly regarding variations between different subjects. We eschewed common movements such as reaching and walking [18, 19, 13] and also studies of small-scale grasping movements [20, 21].

In our study, a full-body virtual-reality tracing task elicited a series of human movement sequences [17]. At each trial, subjects freely chose their initial postures and were given no instructions on how to comport themselves during the tracing process. Participants were tracing three-dimensional space curves at their preferred posture sequences and their postures were continuously recorded using a motion-capture system. Specialized aggregation methods were developed for data analysis that extracted similarities of posture sequences in the face of kinematic variations. The exciting and unsuspected result was that both the movement’s posture sequences and kinematic variations showed striking commonalities across subjects. The obvious inference from the observed similarities of movements across different subjects is that there must be some general principle for humans’ motion commonalities. This regularity of movements across different subjects implies energetic cost should be similar. Moreover, these observations arise from the generally argued principle that the self-selected trajectories should be economical in energetic cost. This argument is reinforced with by progress in the sparse coding of temporal sequences [22, 23] that strongly suggest that trajectories are remembered to obviate the difficulties of computing them online. In addition, if movements are to be stored, the less expensive ones are likely to be preferred[24].

For the energetic cost computation, we took advantage of a forty-eight degrees of freedom dynamic computational model capable of simulating, analyzing, and synthesizing humanoid movements [25]. The model consists of twenty-one body components connected by twenty joints and incorporates several novel features. One innovation is that the joint connections are not treated as perfectly rigid constraints but rather as very stiff springs that hold body parts together like tendons and muscles. The model allows computing instantaneous power from the product of net joint torque and joint angular velocity. The work performed at each joint was determined by numerically integrating the instantaneous powers over the entire tracing task. In this way, the energetic cost of human motions can be computed given motion capture data.

To test this hypothesis that the minimum energetic cost principle is still held in large-scale complex movements the costs of different subjects’ curve traces were computed and compared to the costs of tracing movements under two different kinds of perturbations. In one, the tracing trajectories were slightly perturbed by shifting positions of a particular body part of the dynamic model a small amount for the duration of the trace. In the other, the original tracing path was displaced in certain small increments prior to the trace. The result of both of these kinds of perturbations was that their means of the energetic cost were higher than those of the original curve. In other words, the energetic cost exhibits a classical U-shape with respect to the different posture sequences, with the minimum of the U-shape curve consistent with the cost of the original posture traces, which our subjects self-selected. These results strongly suggest that that movement is selected on the basis of predicted minimum cost.

## 2. Background

In the past, a common way to address the minimum energetic cost principle was to conduct experiments comparing walking and running with many other strange and unpractised gaits [26, 27]. Nowadays, there are three commonly used methods to study energy optimization.

The most straightforward and frequently used method is to measure the metabolic cost, e.g., subjects breath through a mouthpiece to measure oxygen consumption rates (VO2). For example, subjects were required to walk under different circumstances, and the results showed that the metabolic cost was minimum while subjects walked at the condition which was “comfortable” for them [3, 4, 5, 6, 14, 15, 16]. The advantage of this method is that movements can be related directly to energetic cost, but the measuring apparatus is typically very constraining.

A common way to measure muscle co-activation and stiffness is to use Electromyography (EMG). Huang et. al. huang2012reduction showed that that subjects’ metabolic cost is reduced during the learning process of arm reaching tasks, and their muscle activities and co-activation would parallel changes in metabolic power. However EMG measures just a correlate that needs additional modeling to turn it into a energetic cost.

A third energetic cost method, dynamic modelling, is to build a closed form analytical mechanics-based model and determine if the predicted minimum mechanical cost correlates with people’s kinematic preferences. For example,,[7, 8, 9, 11] use an inverted pendulum model to predict the op-timal step length and compare it with the subjects’ real step length while walking.

All these methods pose obstacles for our calculation of the energetic cost of whole-body tracing movements collected from the VR experiment [17]. These methods are time-consuming, and the required configuration restricts the variety of experiments. For example, the VO2 process does not work for our virtual-reality tracing tasks as subjects need to wear the VR helmet on their head, leaving little space for a mouthpiece. Besides, the EMG method measures muscle co-contraction, which is correlated with energetic cost, rather than calculating the cost. Another possible way is to build a humanoid dynamic model. The method is the best way to imitate human movements, and it is widely used in biomedical engineering due to its compliance with real-world physical rules. However, it has several critical limitations as well: 1) it is too difficult to model and control a complex system, such as a whole human body. 2) it is challenging to represent “kinematic loops”, such as postures that need both feet are on the ground. 3) for large systems, the equations of motion in nested, rotating reference frames become very complex, making them more challenging to approximate well. Due to the complexity and disadvantages of dynamic modeling method for large complex systems, most of studies took advantages of two-dimensional models to study human part-body motions in the sagittal plane.

There are some methods of building a dynamic 2D bipedal robot by modeling the whole-body with a skeleton of rigid segments connected with joints. However, those methods over-simplify human bodies so that they can only study simple single-behavior human movements. The simplest bipedal robot uses three links to represent the torso and two legs in the sagittal plane [28, 29]. Five-link biped robots extend the model using two links to represent each leg [30, 31, 32, 33], while seven-link biped robots further extend it by adding feet to it [34, 35]. Furthermore, those methods have many assumptions while studying human locomotion. For example, most researchers assume that when the swing leg contacts with the ground, an instantaneous exchange of the biped support legs takes place. In this way, the biped locomotion with single foot support can be considered as a successive open loop of kinematic chain from the support point to the free ends, as robot manipulators. Recently, 3D modeling of closed-form nodeling of biped robots [36, 37] has been developed. However, they are still not sophisticated enough compared with a real human body.

In the face of these complex challenges, a major alternate modeling route is to forego the neural level of detail as well as one that features muscles and model more abstract versions of the human system that still use multiple degrees of freedom but summarize muscle effects through joint torques. The computation of the dynamics of such multi-jointed systems recently has also experienced significant advances. The foremost of these, use a kinematic plan to integrate the dynamic equations directly. Several different open source dynamic libraries exist, such as MuJoCo^1^ [38], Bullet^2^, Havok^3^, Open Dynamic Engine(ODE)^4^, and PhysX^5^, but an evaluation by [39] found them roughly comparable in capability, and only MuJoCo has been applied to human modeling.

Our 48 degree of freedoms human dynamic model (HDM)^6^ [25] also based on a direct integration method. It was built on top of ODE which is the most commonly used dynamic library in robotic research area. The model has a singular focus on human movement modeling and uses a unique approach to integrating the dynamic equations. A direct dynamics integration method to extracts torques from human subjects in real-time [40, 41, 42] using a unifying spring constraint formalism.

These toques have two components. The major component is the one determined by the open-loop integration of Newton’s equations. These must be supplemented by a closed-loop set of “residual torques” to achieve accurate balance. This organization models the similar dichotomy in the human system.

At each frame, instantaneous power was computed from the product of the net joint torque and joint angular velocity. The work performed at each joint was determined by numerically integrating the instantaneous powers over the entire tracing task. In this way, given motion capture data, we can compute the mechanical cost without building a humanoid biped robot with motion equations. An extensive validation of this dynamic model appears in [25]. Note that it is common to use mechanical measures of work to indicate the metabolic energy consumption [43]. The “energetic cost” mentioned in the following sections means the mechanical cost.

While doing the virtual tracing experiment, subjects freely chose their starting posture and were given no instructions on how to perform themselves. Therefore, participants were tracing curves at their preferred posture sequences. In other words, they traced curves under the conditions which were “comfortable” for them. According to the previous experiments [3, 4, 5, 6, 14, 15, 16], we can expect that the energetic costs of movements with those trajectories should be a minimum or at least locally minimum. To support our conclusion, the cost of original virtual tracing movements and perturbed movements were computed and compared using the human dynamic model. As expected, the energetic cost always exhibits a U-shape while tracing using different postures sequences, with the minimum of the U-shape curve consistent with the original posture traces, which our subjects self-selected. In this way, we are able to demonstrate the energetic cost of original trajectories is a local minimum. The focus of the method section describes experimental protocol in more detail.

## 3. Results

Using the kinematic curve tracing data from [17], we fitted the dynamic model to each of the eighteen subjects and then had the models trace the nine curves that are shown in Fig. 1. The energy cost of tracing paths showed marked regularities in the following aspects of the data that was subject to the following analysis summary:

**Figure 1:**
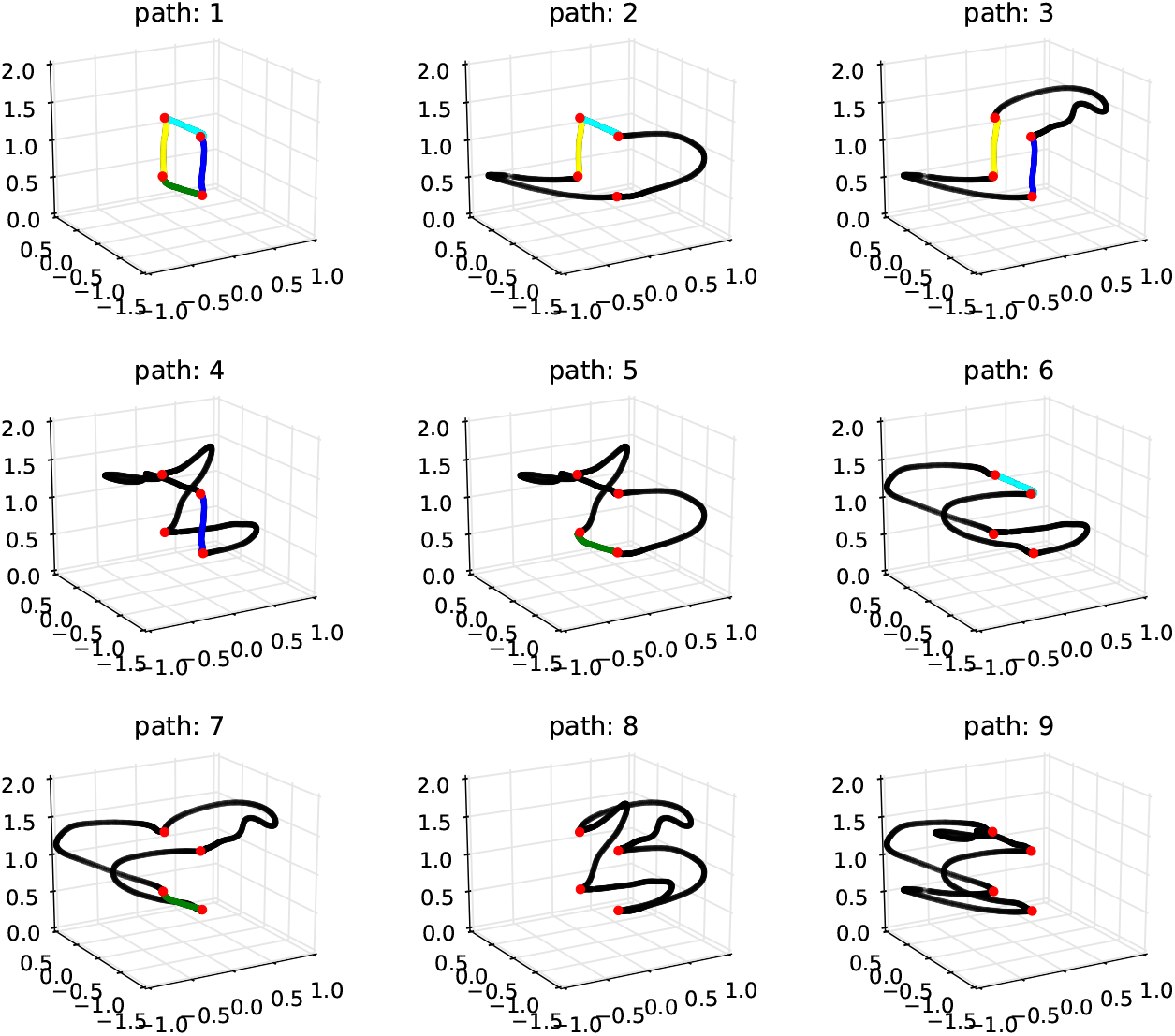
The nine 3-dimensional paths in the virtual environment that were used in the experiment. They are ordered by their complexity. For reference, colors denote common segments and points. For the subjects, the paths were all rendered in black, The scale is in meters.

1. The joints’ power allocation while tracing path 1 across different subjects showed that although the total costs of the movements varied between subjects, the power use is qualitatively very similar. (See section 3.1, Figure 2);
2. The computation of average energy cost while tracing path 1 showed the magnitude of the required residual forces were relatively small. (See section 3.1, Figure 3);
3. The costs of tracing each path by each subject are very similar and approximately monotonic with the length of paths. (See section 3.2 and Figure 4);
4. Although there are variations in the cost across the repeated traces, the cost of using the perturbed model parameters is significantly higher than the original. (See See section 3.2, Figure 5 Figure 6);
5. The increment of energy cost while using perturbed model parameters distributes more on the joints’ cost than on the residual component. (See section 3.2 and Figure 7);

**Figure 2:**
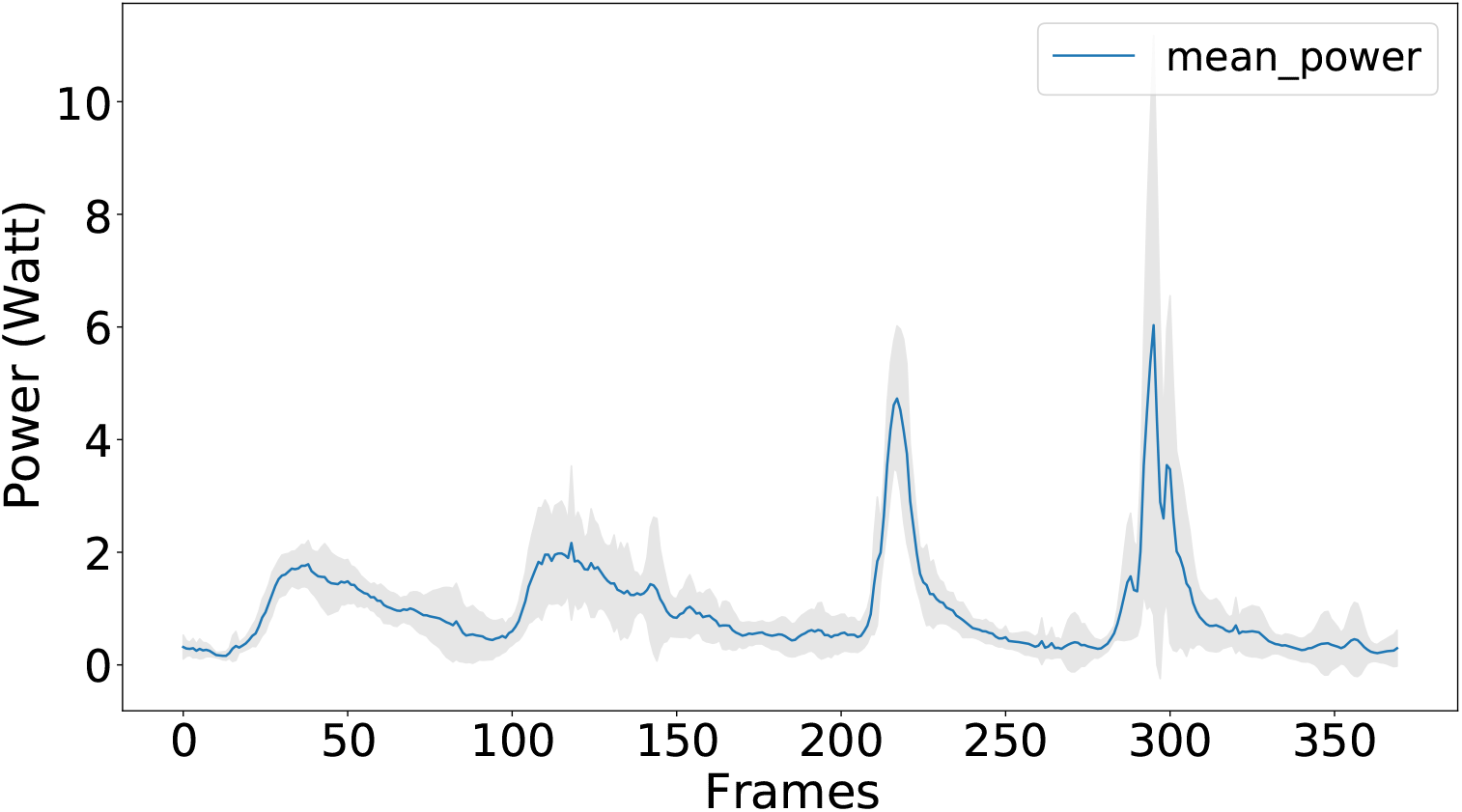
The power of tracing path1 at each frame Nine subjects traced path1 five times. The plot shows the average joints’ power at each frame across subjects. The blue line indicates the mean and the gray shaded area represents the standard deviation of powers. The small standard deviation means that different subjects had similar power patterns while tracing the same curve, which shows that the curve has points of difficulty in tracing shared by the subjects. Path 1 is the most straightforward, but the observation of correlated effort represents patterns in tracing other curves.

**Figure 3:**
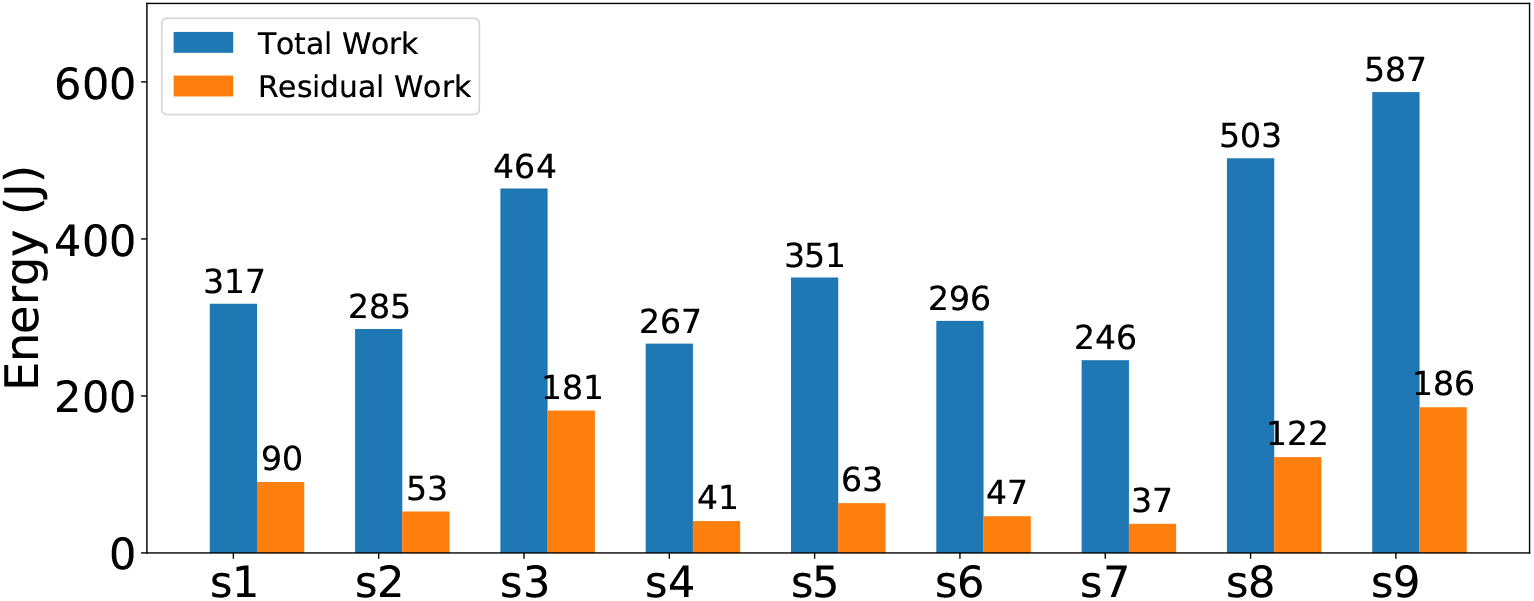
Energetic costs of tracing path 1 Each subject traced path 1 with five repeats. The horizontal labels indicate the related subjects, e.g., “S1” represents the subject1. The total cost is shown in blue, and the portion of that cost due to residual forces are shown in orange. A low cost in residual torque usually signifies that the dynamic model is a good match for that subject’s kinematic data.

**Figure 4:**
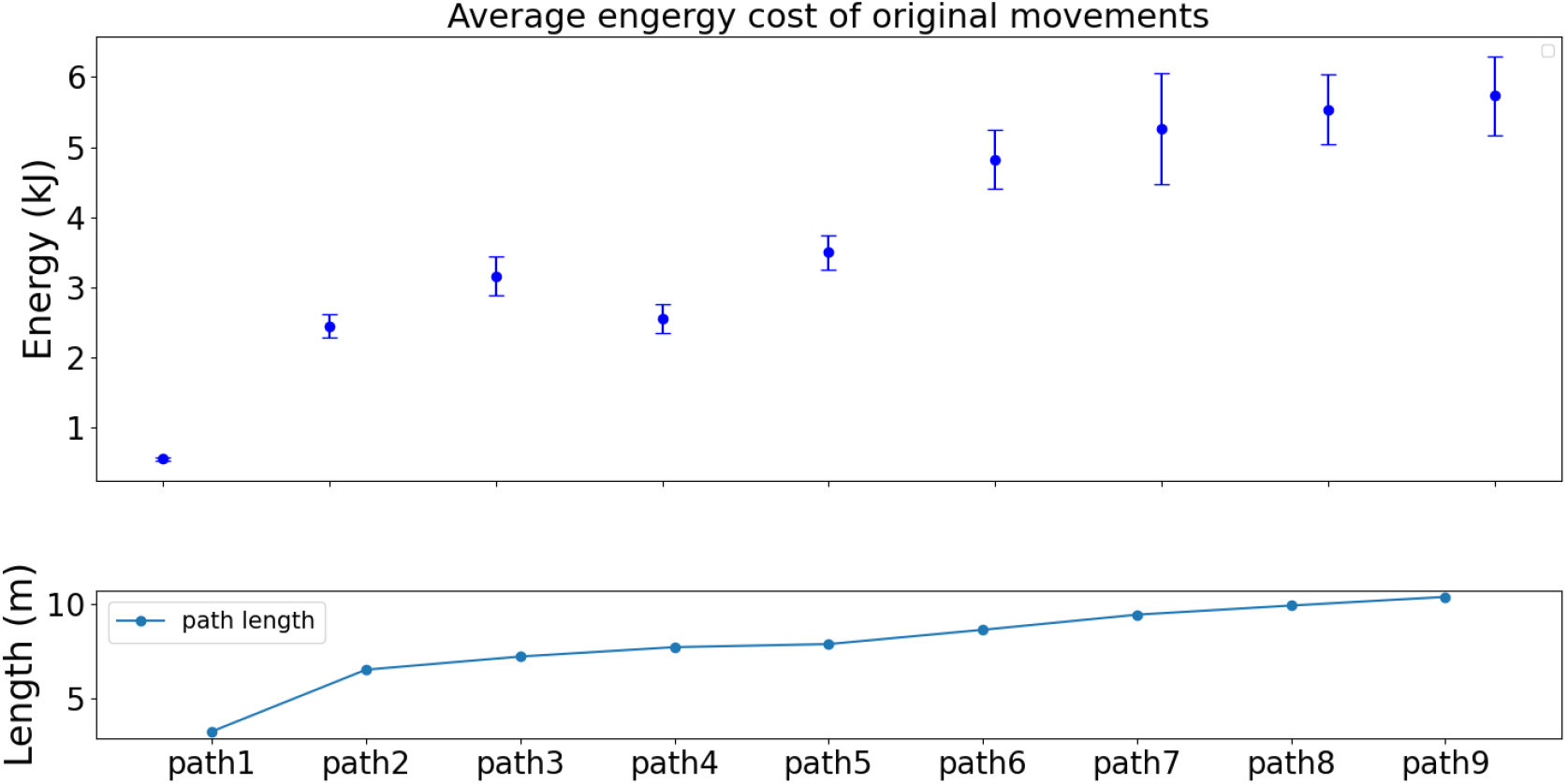
Cost of tracing nine paths These results portray the possibility that the costs vary across the best-fit five subjects. The statistics show that each path traced has a unique cost that distinguishes it from the rest.

**Figure 5:**
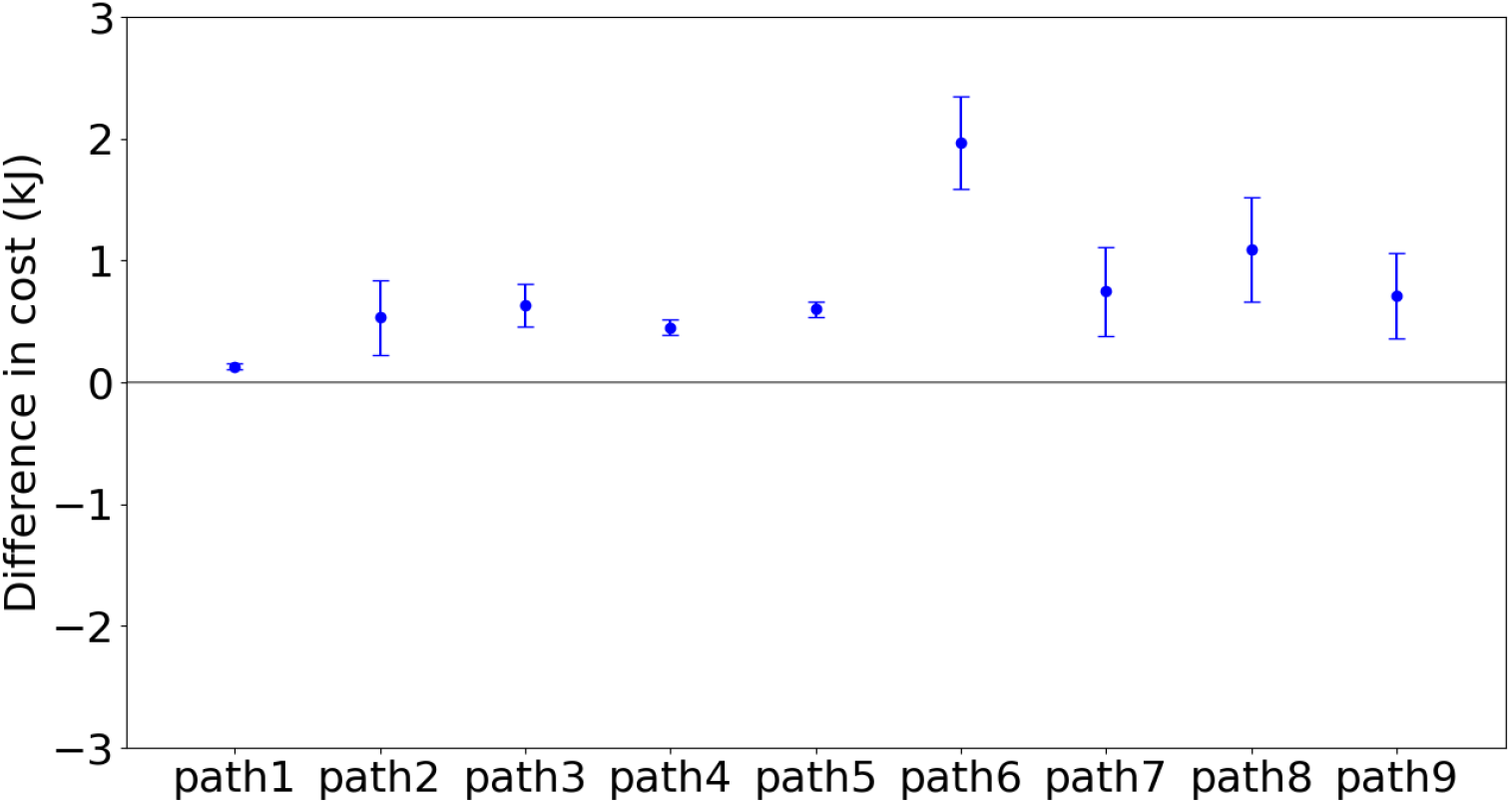
Energetic cost of tracing with model perturbation Energetic cost of tracing each of the nine paths with perturbations in the right elbow marker. The elbow was moved up 5cm. This shows that for all the paths and the averages across subject tracers, the original path is always the least expensive. Moreover, the differences between the energetic costs of original trajectories and perturbed trajectories are highly significant.

**Figure 6:**
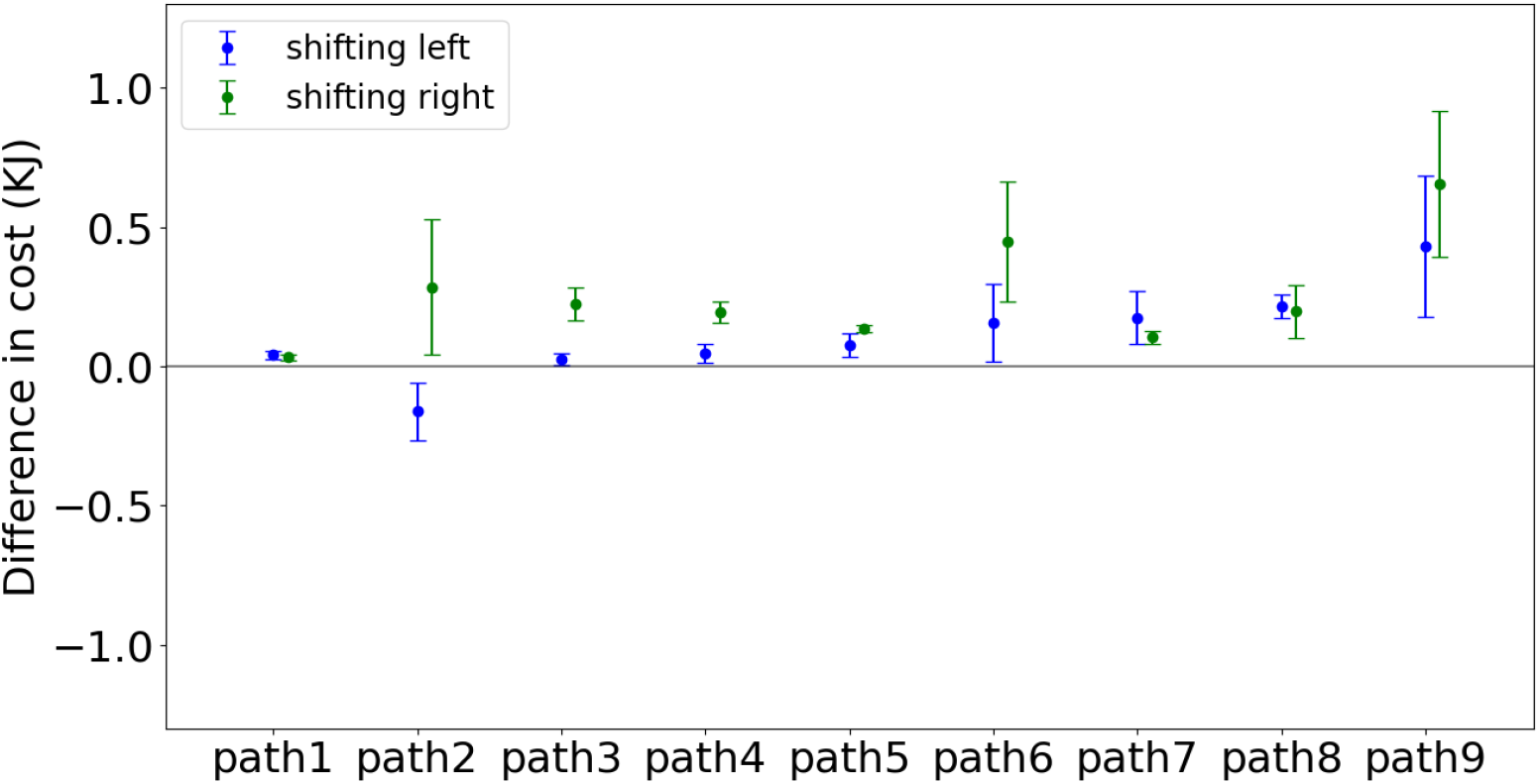
Energetic cost of tracing with path perturbation Each of the nine paths has two perturbations of 5 cm: left in blue, right in green. This main result shows that for both averages across subject traces, the original path is always the least expensive.

**Figure 7:**
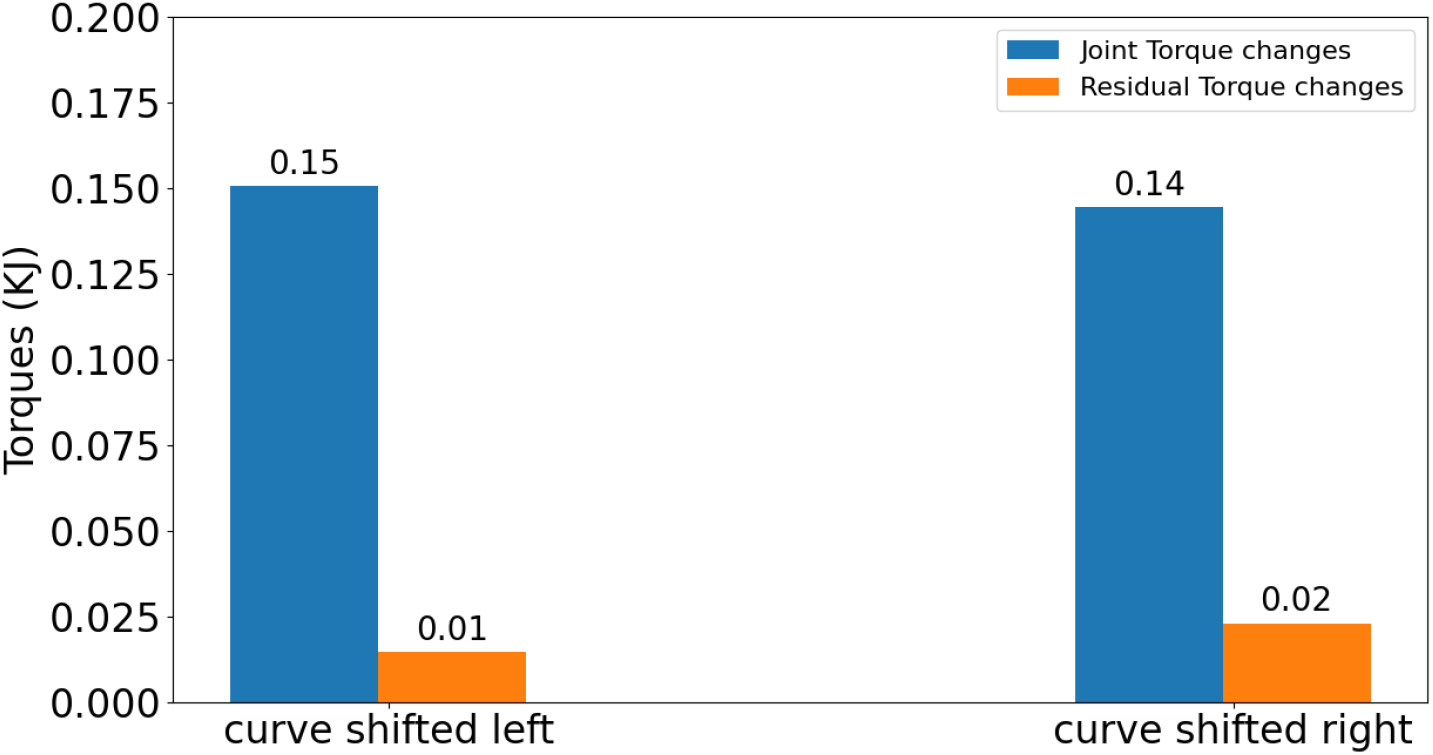
Residual torques The average of the means of the cost changes for path 1 with five repeats across five participants.

### 3.1. Detailed Energetic cost analysis of tracing path1

#### The mean of power across different participants

As an initial analysis, we established the variations in the energetic costs for tracing path 1 exhibited by different subjects. Fig. 2 illustrates the mean and the standard deviation of powers across subjects at each frame. The result reveals that subjects put similar effort at the same points along the path. Thus although the total cost of the movements may vary between subjects, the power patterns are qualitatively very similar. The VR experiment [17] showed participants used similar postures sequences while tracing the same curves from a kinematic perspective. It is expected that the instantaneous power of joints at each frame should be similar as well due to the skeleton constraints of the human body. The similarity of power patterns across different subjects reinforces this conclusion from a dynamics perspective.

#### Average energy cost of five repetitions

Although there are qualitative similarities in the difficult points on the curve, the total costs of the traces differ across different subjects. This result is expected due to the variety of subjects’ skeletons and weights. Fig. 3 represents the energetic cost per subject. The total energy of tracing a path1, including the residual components, is shown in blue, and the residual component is shown separately in orange. When reporting the energetic costs of the traces, we always use the total cost shown here in blue.

#### Residual forces

As shown in Fig. 3, the highest cost of the tracing movement is the component owing to the joint torques that are producing the kinetic trajectories, and the additional cost of the residual from the inverse dynamic calculation is small. In the human system, this residual is most prominently due to the vestibular system, but just how the vestibular connects to the muscular system is not modeled by the human dynamic model. Instead, we implemented a provisional system of torques referred to as a co-ordinate system positioned and the center of mass to maintain balance [25].

### 3.2. Energy cost analysis of tracing individual paths

#### Energy cost of tracing nine paths

Although there are similar energetic costs per subject in tracing a same path, this arrangement does not carry over to the comparison between paths, which has larger differences. We hypothesized that the cost should scale as the length of the path, as shown in Fig. 4, which shows the average energetic cost of tracing the nine different paths. The paths differ in tracing cost, but the costs of tracing each path by each subject are very similar and approximately monotonic with the length of the paths.

Given these regularities, the next step was to evaluate the significance of perturbations in the tracing protocol. The hypothesis is that if the tracing postures are chosen to be of minimum energy, changing the configuration away from the original tracing situation should incur a cost, which was what happened.

#### Model perturbation

The first perturbation test changed in model marker trajectories, called model perturbation. Specifically, the right elbow marker was shifted by a small delta, which produced a new constraint that the model needed to satisfy while tracing paths. To implement it, the dynamic model had to trace paths using the same posture sequences except for lifting its right elbow. Although kinematics of the body parts except the right elbow remained for the unperturbed trace – only the kinematics of the right elbow changed, the joints’ constraints bias the dynamic model adapt to follow the new perturbed trace.

For each trace, the right elbow marker was raised by 5 cm. The rest of the system adapted the way dictated by the dynamic constraints. Fig. 5 shows the difference in cost of constrained motions and original motions. It is seen that although there are variations in the cost across the repeated traces, the cost of using the perturbed model is higher than the original. Note that outside of the changes, the rest of the model solves the inverse dynamic model with the unperturbed parameters, and thus the model has substantial degrees of freedom at its proposal. The significant test showed the difference is reliable, with a p-value less than 0.001. Furthermore, it is obvious that the increase of tracing complex paths is larger than that of tracing simple paths.

#### Path perturbation

The second perturbation test made adjustments in the traced path, called path perturbation. Some effects of displacement can be intuited. For example, if a subject has to reach over their head during the trace, it can be expected that lowering the traced path would result in cost savings. For this reason, we chose path perturbations in the horizontal plane. Two such perturbations were used: a 5-centimeter leftward displacement and a 5-centimeter rightward displacement. Left and right are referenced to the coordinate system used for the four points used for all nine curves (See Fig 1).

In this way, new constraints were produced as the dynamic model was required to trace the perturbed paths while the starting tracing positions were not changed. In contrast to the model perturbation, the model’s trace paths were shifted while the posture sequences remain the same.Again, the dynamic model took advantage of internal joint constraints to adjust original posture sequences to trace the perturbed paths.

Figure 6 shows the difference in average energetic costs for tracing displaced paths and original paths across subjects. The blue dots indicate the difference between motions of tracing left-shifted paths and motions of tracing the original path while the green dots represent the other case. For most cases, the original paths are seen to be consistent with the lowest cost. The path 2 with 5cm leftward displacement costs less than the original path 2. The reason is that subjects preferred to stand near the left corner which is the starting tracing point. However, the left part of path 2 is much easier than its right part (See Fig. 1). Therefore, when shifting the path 2 to left, subjects became closer to the right part, which led to an easier tracing. In contrast, subjects had to move their bodies more in order to trace properly when shifting path 2 to right.

Here again, the overall result is striking. Although there are some overlaps, the original paths are more economical for almost all curves than the displacements. The significant test showed the effects of shitting paths is not very clear but still reliable, with a p-value less than 0.01. The observation that the averages of all the perturbed costs are larger than the average cost of their original progenitors strongly suggests that energy cost is the factor in the choice of tracing postures.

#### Residual forces

Given the dynamics dichotomy, a natural question that arises concerns the magnitude of the extra torques in the perturbation cases. Are the extra costs carried by the dynamic model or the residual? It can be answered by interrogating the simulation, and it turns out that the dynamics model’s contribution is dominating. This is shown in Fig 7.

Note that if the constraints on the dynamics were extremely stiff, then the model would have no course other than tracing an exact copy of the unperturbed trajectory and let the residual torques contribute the needed difference. However, the markers on the body for these experiments were limited to 15∼18 of key body segments, leaving the extra degrees of freedom to be determined by the dynamics. Moreover, the torque computation, to model the reality of muscles [44], used spring constraints at each joint degree of freedom. Finally, the right finger was required to contact the displaced paths, and the remaining features of the movement are the same, leaving the dynamics to fill in the rest.

## Discussion

Given that the cost of the movements is a significant fraction of a human’s caloric budget [45], one might expect that humans would exhibit common low-cost postures. It turns out to be the case for stereotypical situations such as reaching or walking on a planar surface, but arbitrary whole-body movements have been less studied, so the expectations are open. Thus it was a surprise to measure arbitrary movements in a large-scale tracing task and find markedly common posture sequences used by all tested subjects [17]. An obvious possibility for similar posture sequences is energetic cost, especially since there were no complex constraints in the movements and no constraints in the time to perform the traces. Our simulation extends the kinematic finding to show that tests of human dynamics provide evidence that movements are chosen on the basis of energetic economic costs. The cost of tracing scales monotonically with the length of a traced path as expected, and the necessary residual forces, as would be expected from the human’s vestibular system and others, were relatively small, given that the subjects had to choose their movements.

The main substantive results are that subjects’ traces of each of nine space paths all have minimal costs with respect to local perturbations. One manipulation introduced perturbations in their kinematic variables – the subjects traced the path but their model with small displacements in kinematic markers. The other experiment used local horizontal displacements of the paths. Verticals were not used as they can be equivocal. The displacements can interact with the different body heights, e.g., a short subject has to reach an uncomfortable height. However, outside of this caveat, all the data can be interpreted as the tracing posture sequences selected based on energetic cost.

The hypothesis that humans use minimum cost movement trajectories is shown by the use of a human dynamic model that leverages a major innovation in dynamics computation that allows the recovery of torques from kinematic data. The disadvantage of the current method is that we perturbed motions manually, so it is possible that we found only a local minimum in the space of possible movements. However, as tracing a path usually takes more than 1000 frames and at each frame, there are 50 markers representing a posture, the perturbation space is significantly vast. Therefore, our future work is to introduce an algorithm with the capability of seeking potential perturbations automatically, such as reinforcement learning, while still reflecting the constraints of possible postures.

## 4. Methods

### 4.1. Virtual tracing experiment

The original kinematic data capture were collected from a virtual whole-body tracing experiment that was to elicit natural movements under common goals [17]. Subjects wore a virtual-reality helmet, Oculus Rift [46], to see a virtual three dimensional interior room with a dojo backdrop via stereo video. They were required to trace a series of paths positioned at fixed locations in the virtual environment. The movements of their bodies and variables relevant to the tasks were simultaneously recorded using the PhaseSpace motion capture system [47]. The WorldViz Vizard software package [48] both controlled the virtual tracing protocol and the recording of the motion capture data. Fig. 8 shows the virtual environment setup. Fig. 1 shows the nine paths that subjects traced.

**Figure 8:**
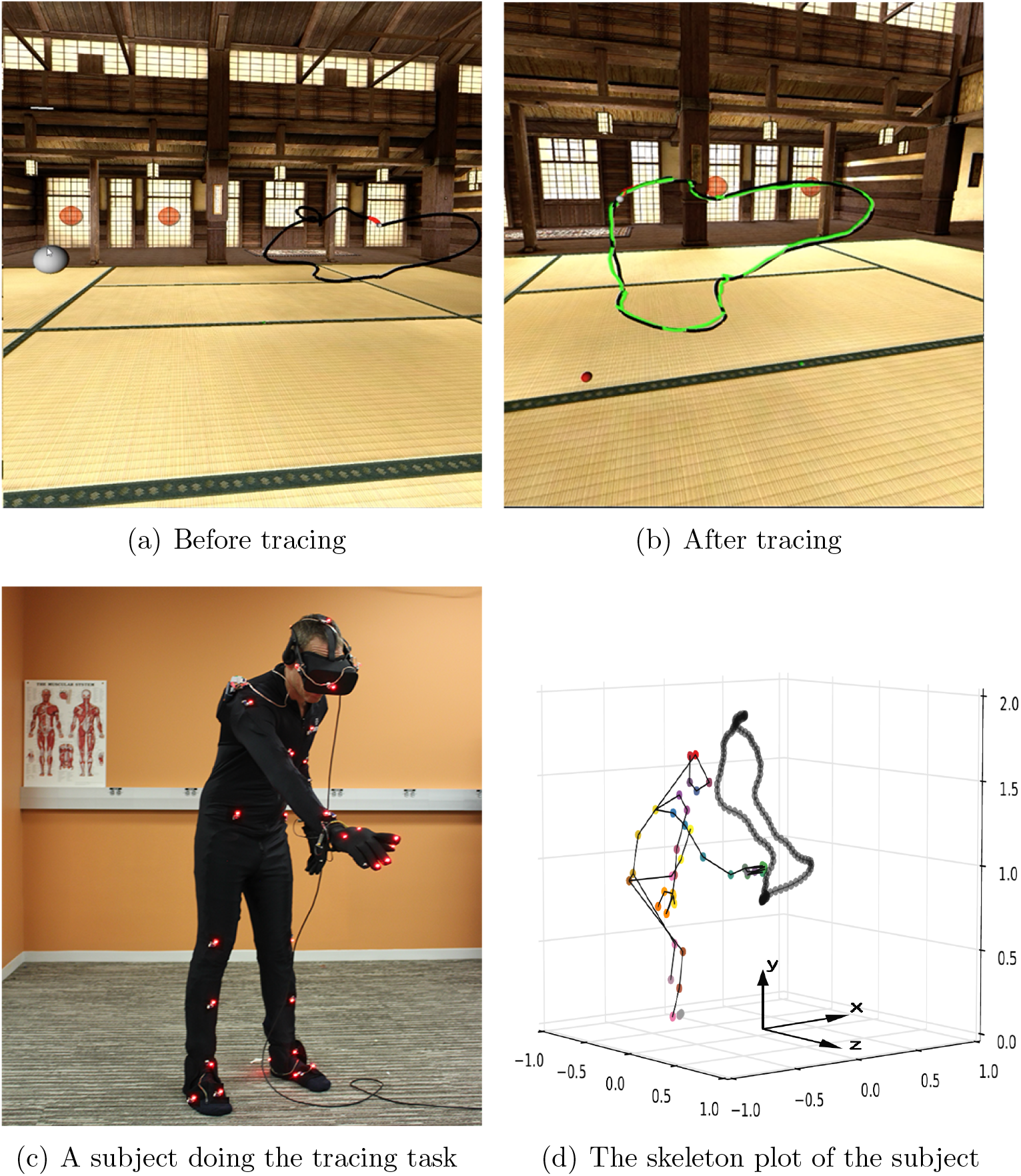
the virtual environment setup. (a) shows a full view of a path, denoted by a black path, and the starting position, denoted by a large white sphere. The small white sphere on the path at the end of a red segment is the tracing target sphere. (b) depicts the scene when a trial is finished. The green path is the actual tracing trajectory generated by a subject. (c) illustrates a subject in the act of tracing a path in the laboratory’s motion capture 2 x 2 x 2 meter volume. and (d) shows the lab coordinate system. The scale on the graph is in meters. The the subject’s skeleton and the traced path in the 3D space are plotted. The color dots correspond to a subset of the fifty active-pulse LED markers on the suit and the virtual-reality helmet.

#### Data pos-processing

For some frames the motion capture system is unable to determine the 3-dimensional location of some markers, thus raw motion capture data usually contains some segments of signal loss (dropouts). Dropouts are relatively infrequent in practice but can occur over significant temporal intervals, which makes linear interpolation a poor choice for reconstructing the raw motion capture data. In this experiment, trajectory-based singular value threshold was implemented to reconstruct missing marker data with a minimal impact on its statistical structure. The data for each subject was interpolated using a separate matrix completion model.

In addition to the data interpolation process, if a participant did not trace the path successfully we would consider this tracing invalid and the data unusable. Because if a recording of a tracing trial failed, e.g., too many markers were off during a tracing, it will lead to extremely large joint torques, which is unrealistic.

### 4.2. Human dynamic model

#### Model topology

To compute the energy cost of subjects tracing paths, we used our human dynamic model [25]. By replaying the virtual tracing experiment’s kinematic data, we can compute can the joints’ properties, e.g. torques and angles, at frame rates. The human dynamic model is built on top of the ODE physics engine [49]. It consists of a collection of rigid bodies connected by joint. Each joint connects two rigid bodies with anchor points (center of rotation) defined in the reference frame of both bodies. Fig. 9 shows the number of body segments and topology of the human dynamic model.

**Figure 9:**
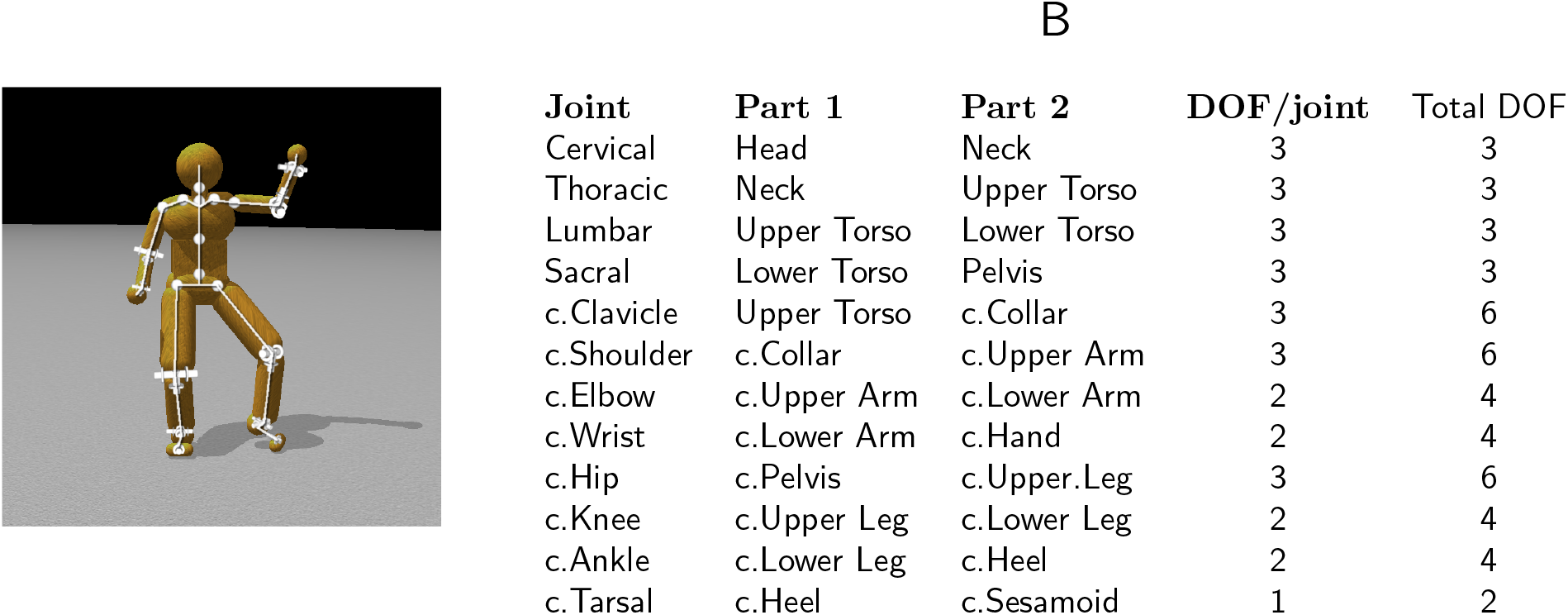
The 48 internal DOFModel. A. Four ball-and-socket joints connect five body-segments along the spine from the head to the waist. Ball-and-socket joints are also used at the collar-bone, shoulder, and hip. B. A summary of the joints used in the model. c. = chiral: there are two of each of these joints (left and right). Universal joints are used at the elbows, wrists, knees, and ankles. Hinge joints connect the toes to the heels. All joints limit the range of motion to angles plausible for human movement. Our model assumes that joint DOFs summarize the effects of component muscles.

Fig. 10 shows a user interface that allows the simulation of human movements via a multi-purpose graphical interface for analyzing movement data captured through interaction with the virtual environment. With this tool, it is possible to interactively fit a model to motion capture data, dynamically adjust parameters to test different effects, and visualize the results of kinematic and dynamic analysis, such as the example in Fig 11, which shows a four stages in a tracing sequence made originally by a participant of the virtual tracing experiment and recreated by applying the inverse dynamics method using this tool.

**Figure 10:**
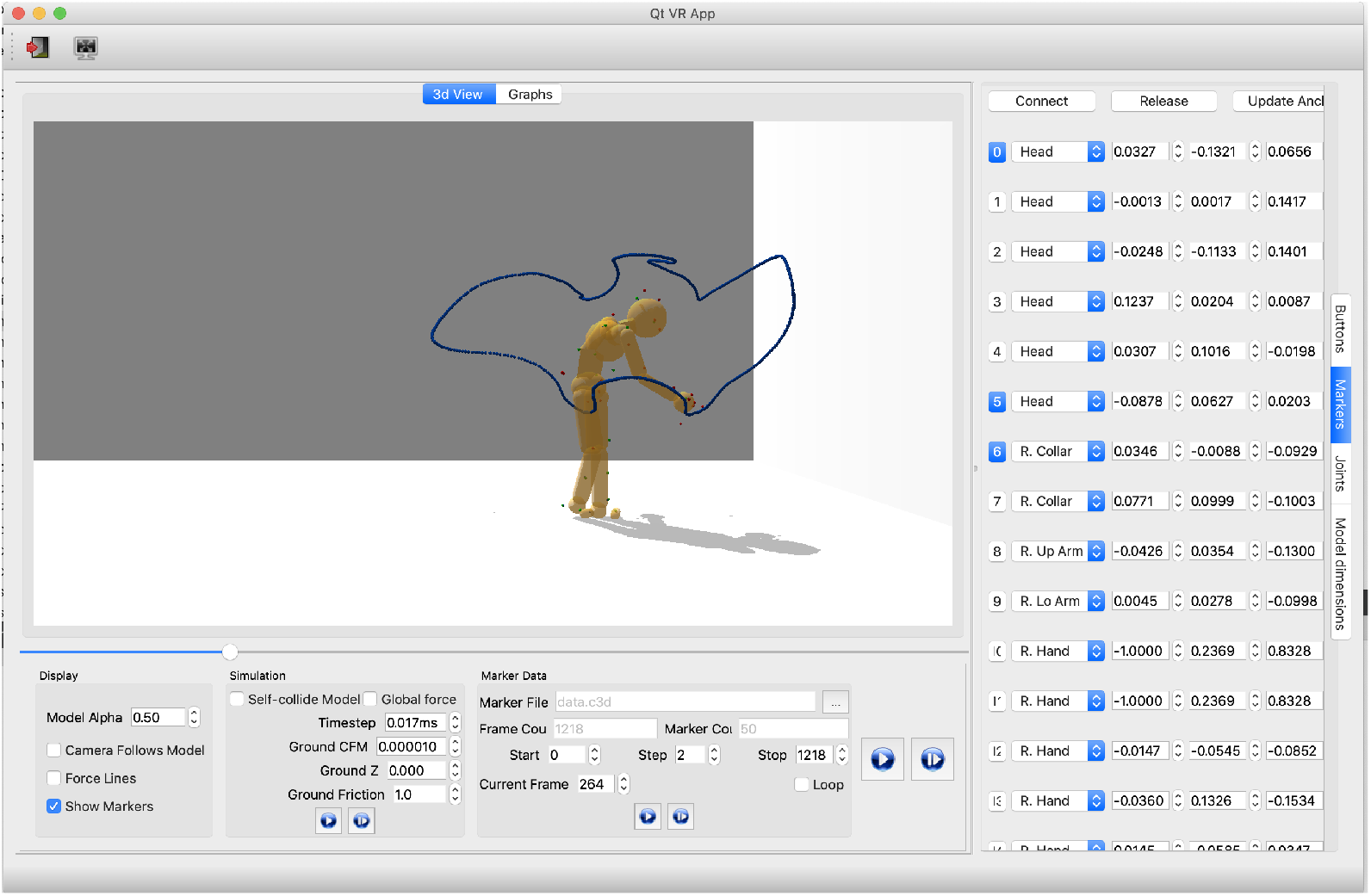
Our analysis tools use the physics engine to compute inverse kinematics and inverse dynamics. They also support various visualizations of relevant data and control for analyzing and producing physically-based movements. The programmed parameters of the model consist of its joints and its 3D marker positions. For example, the right column represents the positions of the markers relative to their corresponding body segments, e.g. the first row shows the information of marker1: 1) “1” represents the marker index, 2) “head” means marker 1 is attaching to the “head” body segment, 3) the remaining three float numbers are marker1’s relative position.

**Figure 11:**
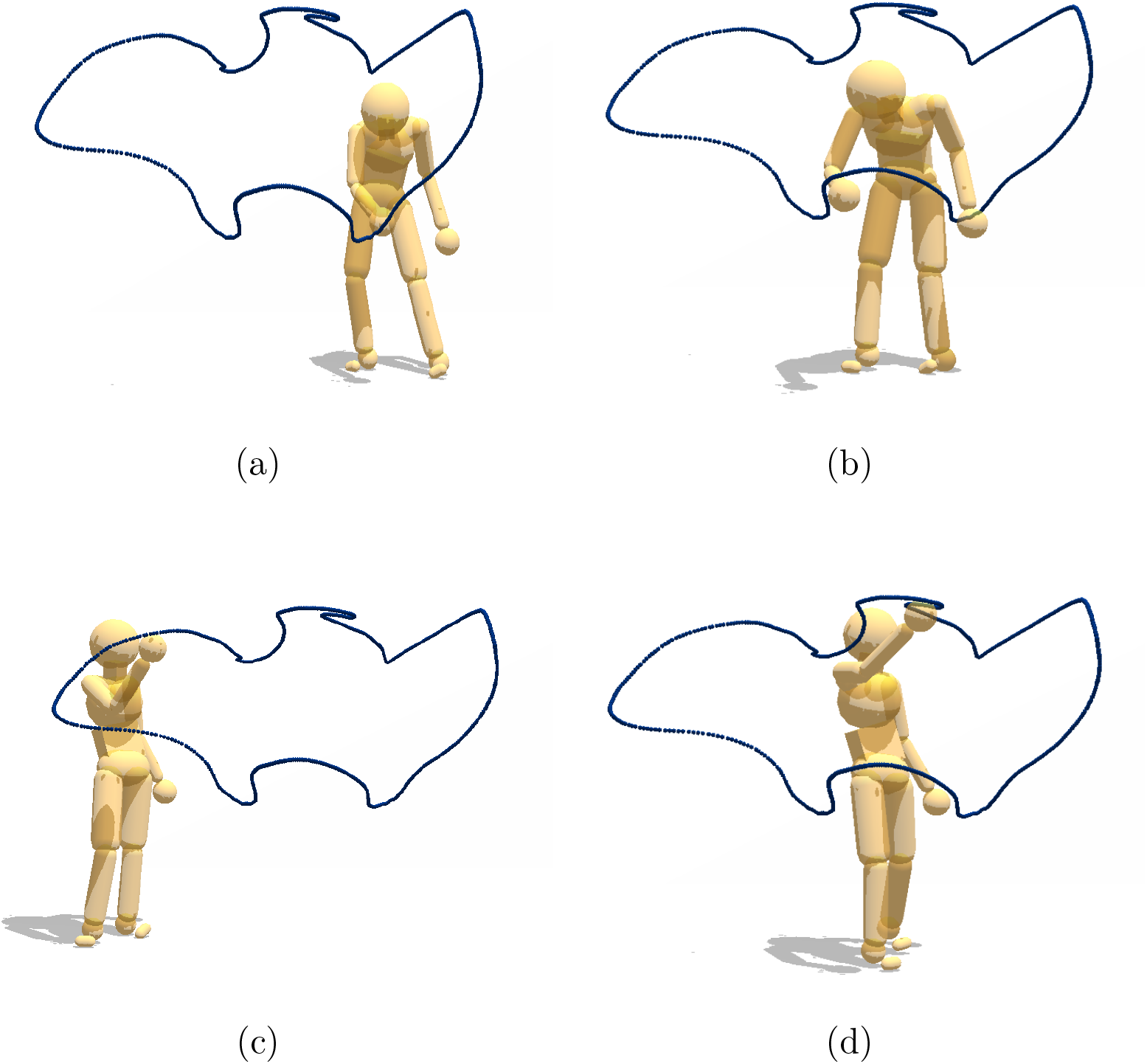
Model capability illustration. Four points in a tracing sequence reproduced with physics-engine-based inverse dynamics using recorded motion capture data from a human subject.

#### Residual forces/torques

The energetic costs are derived from the inverse dynamics technique described in [25], which combines measured kinematics and external forces to calculate net joint torques in a rigid body linked segment model. A feature of the dynamic method is that it can reduce potential errors, both in the matches of the motion capture suit and the model. Analogous to the human body’s ligament structure to join joints, some leeway is allowed in the model joints in the integration process. Nonetheless, even after these adjustments, some errors remain. In the model, the main source of the residual forces is usually attributable inaccuracies in the matches between the motion capture suit makers and their match with their corresponding points on the model. This is commonly resolved by introducing ‘residual forces,’ which compensate for this problem [50]. This resolution with a dichotomy of forces is analogous to the human system, which combines feedforward lateral pathway forces with medial pathway feedback forces. Therefore, a low cost in residual forces usually implies that the dynamic model is a good match for that subject’s kinematic data.

### 4.3. Energy cost computation

The centerpiece of the analysis depends critically on the definition of a posture. At each frame, posture is defined as a vector of the joint torques and angles of each of *N* joints (*N* = 22 in our dynamic human model). The posture *p* at a frame is a 6n-dimensional column vector presenting the joints properties of the *i th* participant, thus

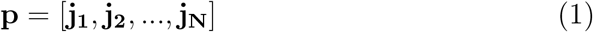

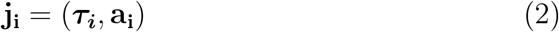

where 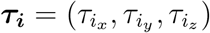 and 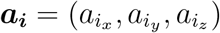 represents the torques and angles of the *i th* joint at a frame respectively and *i* = 1, 2, …, *N*. For the joints which have less than three dimensions, e.g. hinge joints, universal Joints, the values at unused dimension were assigned zero.

The power *W* of *i*_*t*_*h* joint at a frame *t* is a scale and equals to the inner product of its torque *τ*_*i*_ and its angular velocity *ω*_*i*_, thus

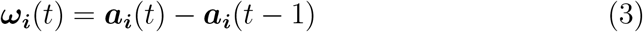

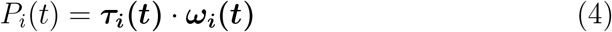

Therefore the power of a posture at frame t is presented as:

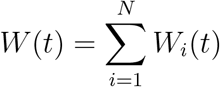

Assuming it takes a participant T frames to trace a path, then the total energy cost *E* of the participant tracing a path is:

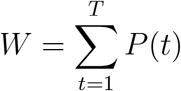

The energy cost analysis is naturally organized into three separate stages. Initially, we analyze the subjects energy cost and residual torques of tracing path1 which is the simplest path. Next, we computed the tracing cost of all nine paths. To compare the energy cost of tracing a path across subjects, we computed the average energy cost for all five repeated traces of each subject. Finally, we measured the tracing cost of perturbed participant’s trajectories and perturbed paths.

## Acknowledgments

This research was supported by National Science Foundation grant CNS1446578 and National Institutes of Health R01 RR09283.

## Footnotes

1 MuJoCo http://www.mujoco.org/

2 Bullet https://pybullet.org/

3 Havok https://www.havok.com/

4 OpenDE: http://www.ode.org/

5 PhysX: https://developer.nvidia.com/gameworks-physx-overview

6 The HDM model: https://github.com/EmbodiedCognition/HDM_UI

